# Ultrastructural features of bacterial calcite unveil ubiquitous organics occlusion and clues on biomineralization

**DOI:** 10.1101/2025.04.15.648918

**Authors:** Franco Grosso Giordano, Martín Purino, Nico Boon, Nele De Belie, Carlos Rodriguez-Navarro

## Abstract

Microbially-induced calcium carbonate precipitation is a widespread natural phenomenon with numerous technical applications. Recent advances have shown that bacterial calcium carbonates (BCC) form non-classically via amorphous calcium carbonate (ACC) precursors in the presence of organics, but the role of organics in the formation and nanostructural features of BCCs is not fully understood. Here we show that two bacterial strains produce BCCs with diverse textural and structural features at the macroscale but similar at the micro and nanoscale. We show that bacterial organics guide precipitation of calcite, stabilizing ACC to produce nanogranular crystals and these organics are then trapped within the crystal, rather than being released as previously suggested. These organics are N-rich and create regions of low Z-contrast aligned perpendicular to the *c*-axis of the crystal, yielding a “Swiss cheese-like” mesostructure. Moreover, it is these occluded organics that lead to the distinctive biosignatures observed in BCC. Finally, we also observe crystalline 2D-films, possibly proteins, templating the oriented crystallization of calcite. These ultrastructural features help to disclose how microbial CaCO_3_ biomineralization takes place leading to improved technical applications and may provide fingerprints for their identification in nature.

## Introduction

Biomineralization is a widespread, cross-kingdom phenomenon whereby organisms induce or control the formation of a variety of minerals in or around them ^1^. A widely occurring biomineralization process is the precipitation of calcium carbonate. It forms the basis of many current biominerals structures – including shells of molluscs, the cell wall of coralline algae, and the cystolith of plants ^1–3^ – and ancient – it is the main mineral fraction in microbialites, some of the oldest fossils in the world ^4^. The bioprecipitation of calcium carbonate is also a common occurrence in the present day microbial world ^5–7^, usually denominated as microbially-induced carbonate precipitation (MICP). While calcium carbonate biominerals in eukaryotes is considered to be tightly controlled by genetic and biochemical factors, MICP, generally an extracellular phenomenon, has long been considered uncontrolled with no specific purpose or functionality ^8^. However, recent evidence is questioning this notion ^9^. For example, calcium carbonates have been reported to lead to an evolutionary benefit by increasing the pathogenic capacity of strains ^10^, facilitating biofilm formation ^11^, and providing survival strategies in extreme conditions ^12^.

Certain aspects of the biochemical understanding of MICP are well agreed upon. For instance, the metabolic pathways of bacterial calcium carbonate (BCC) production are in general well-known and include sulphate reduction ^13^, photosynthesis ^2^, denitrification ^14^ ureolysis and ammonification ^15^ and metabolism of organic acids ^16,17^. They all result in alkalinization, thereby fostering the formation of CO_3_^2−^ ions and enabling precipitation of CaCO_3_ in the presence of Ca^2+^ ions. Moreover, researchers recognize that there is a role of extracellular polymeric substances (EPS) - which include polysaccharides, proteins and nucleic acids - in the formation of precipitates, acting as a Ca^2+^ adsorption template ^18^. Besides their role in concentrating Ca^2+^, there is growing evidence showing that the organic-mediated formation of amorphous calcium carbonate (ACC) by prokaryotes prior to crystallization of anhydrous CaCO_3_ crystals appears to be a general phenomenon ^19–22^. Recently, Enyedi et al. ^23^, showed that bacterial EPS led to long-term stabilization of ACC, and suggested that the stabilized ACC eventually led to the formation of calcite through non-classical crystallization. However, the MICP crystallization mechanisms are still poorly understood, specifically when the role of organics is involved, while structural aspects of these biocrystals have not been fully characterised. This is an important research gap given MICP occurs with a wide range of organisms and it has attracted a large amount of research in bio-engineering applications ^7^.

In this work, we aim to shed light on the formation mechanism of MICP and study the elemental, compositional and (ultra)structural characteristics of BCCs at multiple scales. To achieve this goal we have performed crystallographic, compositional, textural and (nano)structural analyses of BCCs (calcite) from two bacterial species, using inorganic calcite as control. Differences between BCCs and inorganic calcite were starker than between BCCs of the two species and were attributed to the presence of (N-rich) organic biosignatures. Furthermore, BCCs formed through non-classical ACC-to-calcite precipitation and produced mesocrystals with a Swiss-cheese like structure with regions of occluded intracrystalline organic matter. In addition, a bacterially-derived 2D crystalline film was identified which enabled the oriented nucleation of bacterial calcite, supporting the role of organic molecules in BCC formation.

## Materials and Methods

### Synthetic calcite preparation

Calcite was produced synthetically in order to act as an abiotic control. A reaction pathway was chosen as proxy to that expected in the bacterial cultures, whereby calcite precipitated by the dissolution of CO_2_ from bacterial respiration. A saturated calcium hydroxide solution was prepared by adding 5 g Ca(OH)_2_ to 100 ml of distilled water, stirring for 30 min at 50 RPM and letting sit for 2 h. The supernatant (i.e., a saturated solution of Ca^2+^ and OH^−^ (equation 1) was poured into a clean container. CO_2_ was then purged through this solution. This enabled the fast dissolution and hydration of CO_2_, creating bicarbonate (equation 2) and carbonate ions (equation 3). Subsequently, CaCO_3_ spontaneously precipitated through the reaction of carbonate ions with Ca^2+^ (equation 4) in the absence of any background electrolyte. The precipitates were filtered using a vacuum pump and polycarbonate filters (φ = 0.22 µm). The filter with the precipitates was placed at room temperature (*T*) and dried overnight. The dry solids were stored in plastic vials for further analysis.

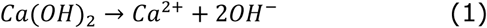

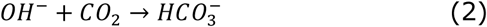

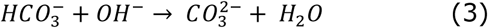

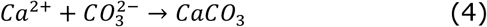

### Selection and cultivation of bacterial strains

Two bacterial strains were used in this study, *Lynsinobacillus fusiformis* (LMG 9816) and a recently isolated strain, *Shouchella clausii* (registered at the Belgian Coordinated Collection of Microorganisms as LMG 33400). *S. clausii* was isolated from a lime mortar wall in Ghent, Belgium by selecting morphologically distinct colonies from a Horikoshi medium. Its capacity for carbonate precipitation was tested on pH 8-adjusted B4-medium (0.4 % yeast extract, 0.5 % dextrose, 0.265 % Ca-acetate hydrate, 1.4 % agar, adjust pH to 8 using 0.1M NaOH), a standard medium used to study precipitation of calcium carbonate crystals ^5,23–25^. Importantly, Ca-acetate was filter sterilized as doing otherwise would lead to precipitation of calcium carbonate either through too high pH or through heating of the Ca-acetate. Finally, the strain was selectively adapted to high pH tolerance by repeated inoculation in TSB adjusted to pH 11 with CAPS buffer. An alkaliphile was desirable as MICP commonly occurs in alkaline environments. The bacteria were grown and kept in TSA at 28 °C.

### Culture and precipitation media, and precipitate extraction

Due to the alkaliphilic nature of both bacteria, carbonate precipitation experiments were carried out on pH 8-adjusted B4-medium. For the inoculation of B4 plates, bacteria were inoculated in TSB and grown for 48 h. Subsequently, 10 µL of a fully grown culture was drop-inoculated onto 9 spots on B4 plates. The cultures were grown for 4 weeks at 28 °C. At this point, precipitates were visible on the plate and were extracted. Briefly, the B4 agar was cut into pieces and placed in a glass container and melted in a microwave oven. 100 mL of warm deionized water were added to the beaker to dilute the melted agar. The dilute agar was centrifuged at 5000 rpm for 1 min while still warm. The pellet was then microwaved again, warm water was added, and centrifuged again. This was performed a third time but centrifuged at 7000 rpm. The sample was warmed up once more, and the suspension was filtered using a vacuum pump and polycarbonate filters (φ = 0.22 μm). The filter with the precipitates was placed at room temperature and dried overnight. Once dried, the solids were placed in a plastic vial for further analysis.

### Bleaching of precipitates

A fraction of the bacterial precipitates was bleached to remove any potential organic matter on their surfaces ^26^. Bleaching was done using commercial bleach and deionized water (1:1 dilution) poured in an Eppendorf tube (1.5 mL) filled with 0.1 g solids, followed by incubation for 2 h in a shaking table. The samples were then centrifuged for 1 min at 5000 RPM and the supernatant discarded. Bleached solids were resuspended in MilliQ water, vortexed for 1 min, and centrifuged again. Washing was performed thrice. The precipitates were dried at 50 °C overnight.

### FTIR spectroscopy

Fourier Transform Infrared Spectroscopy (FTIR) spectra of the precipitate were recorded on a NICOLET 20SXB FTIR (400–4000 cm^−1^> spectral range; resolution of 2 cm^−1^>). Baseline and smoothing corrections to the FTIR spectra was performed using Spectragryph v1.2.16.1.

### X-ray diffraction (XRD)

The crystallographic features of the synthetic (control) and bacterial precipitates were analysed with a Xpert Pro X-ray powder diffractometer (Panalytical, The Netherlands) in reflection mode using copper radiation to determine θ−2θ scans (measurement parameters: Cu Kα radiation λ = 1.5405 Å, 45 kV, 40 mA, 3 to 70° exploration range, with 0.00418° step size and 29.8 s integration time per step). Bacterial precipitates were analysed in zero-background Si sample holders. XRD patterns were analysed using the computer program HighScore Plus Version 3.0d.

### TG analysis

Thermogravimetric analysis (TGA) was performed using a TGA 2 (Mettler-Toledo, Switzerland) with a large furnace and an autosampler, using 70 μL alumina crucibles. Samples (12 mg) were heated at 20 °C/min from 25 to 950 °C under air atmosphere (80 mL/min).

### Synchrotron High-Resolution Powder X-Ray Diffraction (HRXRD)

HRXRD analyses were performed at beamline CRYSTAL of the SOLEIL synchrotron (Paris, France). The wavelength 0.672356 Å (18.44024 keV) was selected with a double-crystal Si (111) monochromator and determined by using Si640d NIST standard (*a* = 5.43123 Å). The CRYSTAL beamline is equipped with a high-throughput position sensitive detector (PSD) MYTHEN, optimal for time-resolved experiments. Borosilicate glass capillaries of 0.3 mm diameter were loaded with powder samples and rotated during data collection to improve diffracting particle statistics. Isochronous annealing (10°C min^−1^; 30 s stabilization time at target *T*, tolerance ± 5°C) was performed using a Cyberstar hot gas blower with a Eurotherm *T* controller (Eurotherm, Worthing, UK), collecting diffraction data each 50°C during the heating ramps (50° to 400°C). Once the maximum *T* was reached, the samples were cooled to room *T*, and a final measurement was taken. The data acquisition time was ∼5 min per pattern, with up to two iterations per measurement to obtain a good signal-to-noise ratio over the 0-55.34 °2θ angular range. To calibrate the equipment and reduce possible shifts during experimental analysis, a LaB6 standard was used. An instrumental resolution factor analysis was performed in each case. Under these data acquisition conditions, the angular resolution was better than 0.006° FWHM. As opposed to conventional XRD, such an angular resolution enables to detect lattice strains of 10^−3^ to 10^−4^ associated with intracrystalline organics.

The Rietveld refinement method ^27^ was used to extract lattice parameters of calcite matching the experimental diffraction peaks with those included in Markgraf and Reeder ^28^ using Topas software. Optical quality (highly pure) geologic calcite (Iceland spar, from Chihuahua, Mexico) was used as standard to compare XRD data from BCC.

To investigate the microstructural variations due to organic (macro)molecule incorporation in the calcite structure, the Thompson-Cox-Hastings pseudo-Voigt function was used to fit the profile of the experimental diffraction patterns. Refinement cycles were performed to ensure reproducibility of each analysis, selecting a Goodness of Fit of <9. Note that, as opposed to BCC, in the case of the Iceland spar control no changes in Bragg peak position or broadening (FWHM) were observed after annealing, i.e., after returning to room *T* following heating up to 400°C. These results show that the selected geogenic calcite is a proper reference to compare the results of BCC HRXRD analysis.

### FESEM and TEM analysis

Field Emission Scanning electron microscopy (FESEM) images were collected using an Auriga (Zeiss, Germany) microscope with field emission gun operated at 5 kV. All samples were carbon coated prior to imaging.

Nanoscale features and elemental composition of the precipitates were studied using TEM on a Titan (FEI) with acceleration voltage of 300 kV. The TEM was equipped with a high angle annular dark field (HAADF) detector for Z-contrast imaging and energy dispersive X-ray spectroscopy (EDS) for microanalysis (performed in STEM mode). Right before TEM analysis the solids were dispersed in ethanol, sonicated for 2 min and fished with carbon-coated TEM Cu grids. Imaging was performed using a 20 μm objective aperture, which is a good compromise for amplitude and phase contrast (high resolution, HRTEM) imaging. Selected area electron diffraction (SAED) patterns were collected from areas ∼200 nm in diameter.

### Solid-state NMR (ssNMR)

The 1D experiments were recorded on a Bruker Ascend 500.13MHz Avance III spectrometer equipped with a Bruker broadband 4 mm 1H/X dual channel CP-MAS smartprobe probe head, equipped with Z-gradients, a MAS III pneumatic unit and running Topspin 3.6.5. For each measurement, the spinning frequency was set to 12000Hz, all measurements were performed at room temperature (25°C) and controlled with a Eurotherm 2000 BVT controller and a BCU-I.

All samples were prepared by filling the 4mm Bruker K1910 ZrO2 rotors with an appropriate amount of dry and finely grained sample material (25-30 mg per sample). Using subsequent rounds of filling and packing the rotor led to a tightly filled rotor sample for each material under study. Following the packing of the rotors, each of them was sealed with a Kel-F insert, a Kel-F rotor cap and visually checked for any leftover voids or air pockets.

The spectra recorded on the samples included 1D ^1^H and 1D ^13^C measurements. For the 1D ^1^H ‘onepulse’ experiments, the spectral width used was 200.00 ppm with 4 scans of 4K data points each being accumulated, preceded by 2 dummy scans and a relaxation delay of 2 seconds. Finally, the spectrometer excitation frequency was set on resonance with the frequency (5 ppm) of the most intense signal. Processing consisted of one order of zero filling to 16K real data points, followed by Fourier transformation, phase correction and a base-line correction.

For ^13^C measurements a ‘hpdec’ measurement type was used for non-cross-polarization and ‘hX.cp’ for measurements with cross-polarization (CP) with ^1^H. The spectral width used was 300.00 ppm with 2048 scans of 4K data points each being accumulated, preceded by 4 dummy scans and a relaxation delay of 5 seconds. Finally, the spectrometer excitation frequency was set on resonance with the frequency (100 ppm) of the most intense signal. Processing consisted of one order of zero filling to 16K real data points, followed by Fourier transformation, phase correction and a base-line correction.

Chemical shifts were indirectly referenced by adapting the magnet ‘field’ value -by measuring an adamantane reference sample-, and adapting the magnet field until one of the adamantane signals is at the right chemical shift value (37.77 ppm). All spectra were processed using TOPSPIN 4.1.3.

## Results

### BCCs have distinctive biosignatures associated with organics and ACC

The calcium carbonate precipitating capacity of two bacterial species was studied. *Lynsinobacillus fusiformis* (LMG 9816) strain was selected as it has been commonly used in other studies of MICP ^29,30^. To test whether nanoscale characteristics between BCCs were similar, a recently isolated strain producing morphologically distinct BCC, *Shouchella clausii* (previously known as *Alkalohalobacillus clausii*) LMG 33400, was used. No previous *Shouchella* (or *Alkalohalobacillus)* strain has been reported to precipitate calcium carbonate.

Optical microscopy imaging (Figure 1a,e) disclosed morphological differences between mineral precipitates produced by *L. fusiformis* and *S. clausii*. *L. fusiformis* mainly produced irregular precipitates with spherical incrustations, while *S. clausii* induced the formation of spherical or oval-shape precipitates. A detailed observation of the latter shows that they display elongated disphenoidal forms, or spindle- and/or dumbbell-shaped structures, as also observed in other bacterial species ^31^. Such diverse precipitate morphologies have been observed for different bacteria in studies using the same standard culture medium used here (B4) ^24^. Both bacteria produced precipitates associated with the bacterial colonies as well as in the surrounding media. Field emission scanning electron microscopy (FESEM) analysis showed the presence of carbonate precipitates closely-associated with organic matter (Figure 1b, f). At the microscale all precipitates had an overall rhombohedral morphology, consistent with the presence of calcite (see XRD results below) and were better developed in the case of *S. clausii* samples (Figure 1d vs g and h). All of them showed a nanogranular texture, implying formation through particle attachment (PA). In PA, ACC particles 100 – 400 nm in size ^32^ in the presence of organics ^33^ or crystalline CaCO_3_ nanoparticles ^34^, attach forming a single crystallographic unit and this is typically observed in carbonate biominerals produced by a range of organisms (e.g., molluscs, barnacles, echinoderms, cnidaria) ^26,35–37^, including other bacteria ^23^. The precipitates were bleached to remove any organics on the precipitate surface and study possible crystallographic and compositional changes. After bleaching, organic matter was not observable anymore on the precipitates (Figure 1c, d and h), calcified bacterial casts could be spotted (Figure 1c), and the crystallographic faces were more obvious (Figure 1h). Note that we performed bleaching tests with increasing reaction time from 10 up to 480 min, observing (using FTIR) that the removable fraction of organics could not be detected after 10 min (see Supporting Information and Supporting Figure 1 and Supporting Figure 2).

**Figure 1.**
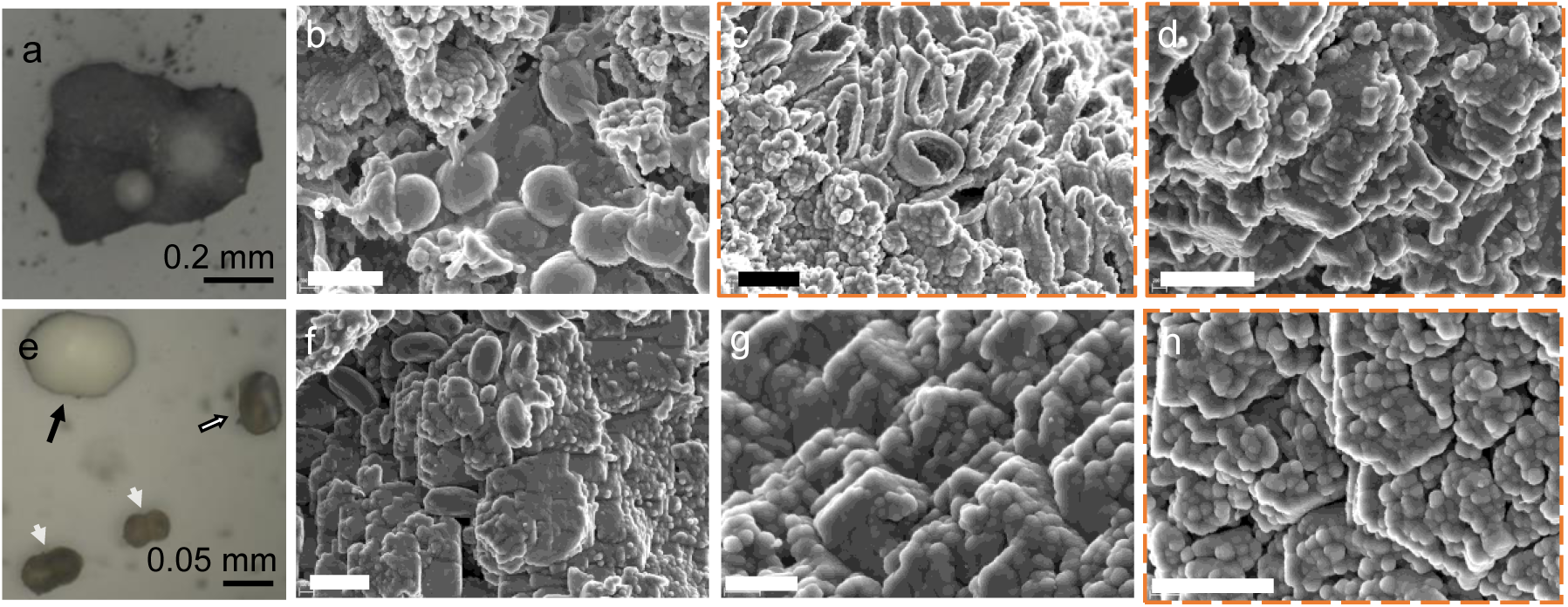
Optical and scanning electron microscopy photomicrographs of bacterial calcium carbonate. Calcium carbonate structures precipitated by *L. fusiformis* (a - d) and *S. clausii* (e - h). Orange outlined images are samples treated with bleach (c, d, h). Different morphologies of calcium carbonate precipitates were produced by both bacteria. (**a**) *L. fusiformis* produced irregular precipitates. (**b**) Bacteria and biofilm associated with precipitated calcium carbonate. (**c**) Bleached *L. fusiformis* precipitates showing the calcified bacterial casts. (**d**) Bleached *L. fusiformis* precipitates (showing rhombohedral vicinal faces) made up of ordered nanoparticles. (**e**) *S. clausii* produced ovoidal (black arrow), disphenoidal (white arrow with black outline) and dumbbell (white arrows) shaped precipitates. (**f**) *S. clausii* cells associated with precipitated calcium carbonate (calcite rhombohedra). (**g**) Detail of the mesostructured CaCO_3_ crystals. (**h**) Bleached calcium carbonate showing the orderly arrangement of calcium carbonate nanoparticles. In all cases the shapeless precipitate subunits varying between 100 – 200 nm in size are distinguishable within calcite mesocrystals. All SEM micrograph scale bars are 1.2 µm.

XRD confirmed the precipitation of calcite rather than any other calcium carbonate polymorphs. The humps at 10 and 20 - 30 °2θ in the XRD patterns of both *S. clausii* and *L. fusiformis* carbonates pointed to the presence of two distinct amorphous phases (Figure 2a). Note that no humps were present in the XRD pattern of abiotic calcite (control). Changes in the relative intensity of Bragg peaks were observed for BCCs, as compared with synthetic calcite. Notably, the intensity of the 110 Bragg reflection increased drastically (as compared to the main 104 Bragg reflection). The (11.0) plane is associated with crystals elongated along the *c*-axis ^38^ and consistent with the crystal morphology observed in Figure 1. Overdevelopment of {11.0} faces, or more exactly, pseudo-faces composed of (01.2) and (11.0) forms ^39^, is common in - although not exclusive to - other calcite biominerals and abiotic calcite formed via ACC-to-calcite crystallization in the presence of organics (see Supporting Information). After bleaching, both humps at 10 °2θ and 20-30 °2θ disappeared. The bleaching of precipitates also led to changes in the intensity of Braggs peaks of both bacterial crystals, most starkly non-(104) planes, with 110 and 113 Bragg reflections being the strongest (Supporting Table 1). We observed better development of crystal faces after bleaching (Figure 1c, d and h) which would explain the XRD changes.

**Figure 2.**
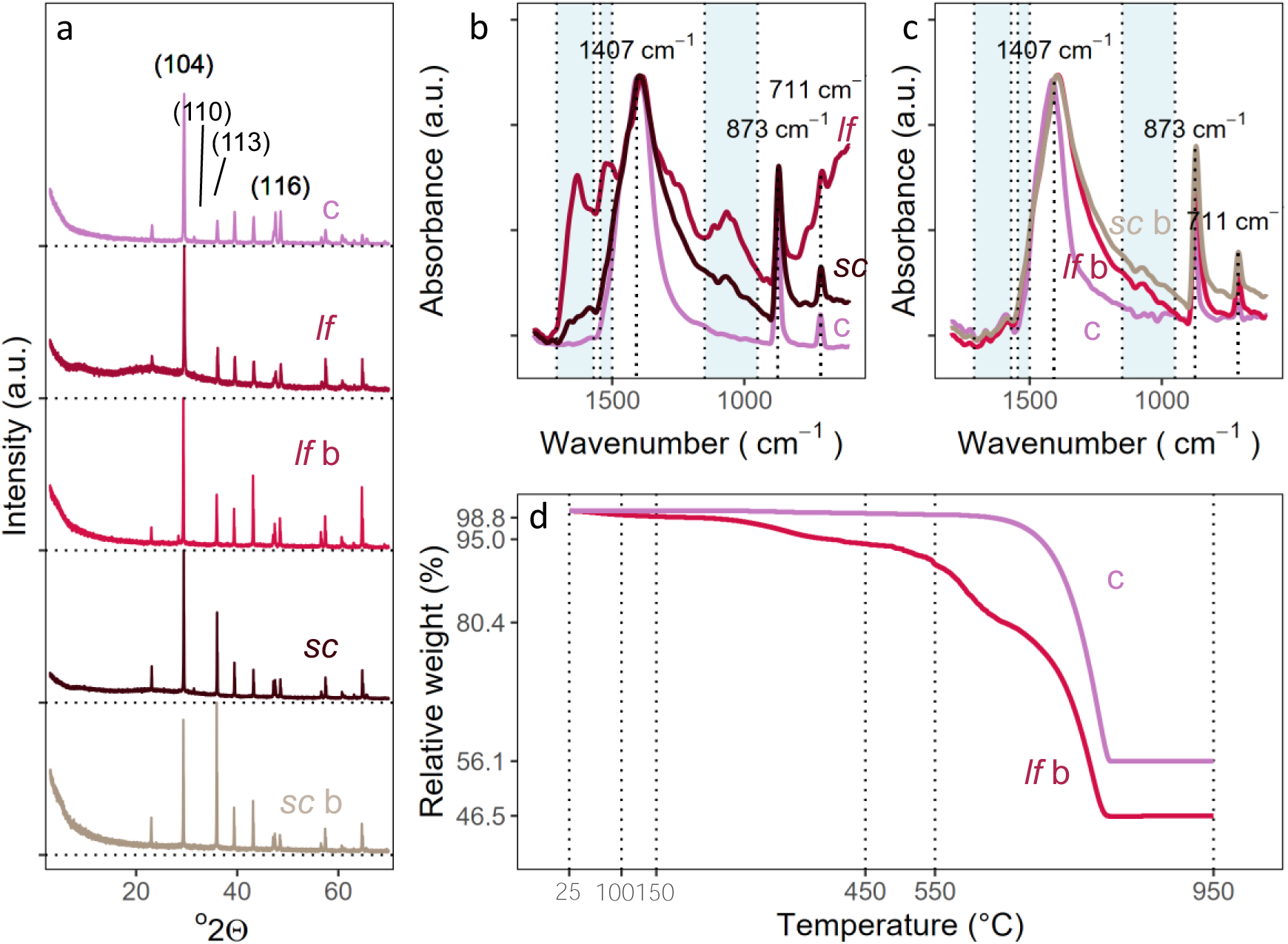
Analysis of synthetic (c, purple) and bacterial crystals (*L. fusiformis* (*lf*, red) and *S. clausii* (*sc,* brown)). Both bacterial calcites were bleached to remove organic matter from the surface (*L. fusiformis* (*lf* b, light red) and *S. clausii* (*sc* b, light brown)) **(a)** XRD patterns of the studied calcium carbonates. Bragg peaks correspond to calcite in all cases and the main *hkl* reflections are indicated. **(b)** FTIR spectra of synthetic, *L. fusifomis* and *S. clausii* calcites. Blue shaded bands indicate organic and/or ACC wavenumber regions. **(c)** FTIR spectra of the calcites after bleaching **(d)** TGA of *L. fusifomis* bleached calcite (light red) vs synthetic calcite (purple).

FTIR spectroscopy showed distinctive spectral features among the different precipitates. All precipitates displayed the characteristic v_2_, v_3_ and v_4_ bands of calcite at 873, 1407 and 711 cm^−1^, respectively, however with some differences. Bacterial calcites showed a slightly red-shifted v_2_ band at 869 cm^−1^ (with a broad tail spanning to lower wavenumbers) and a v_3_ band at 1402 cm^−1^. The red-shift of the v_2_ band can be caused by ACC which has a characteristic band at 860 cm^−1^ also seen in ACC producing bacterial strains ^3,40–43^. There were also variations in the absorbance of the v_2_ and v_4_ bands. Synthetic calcite had the sharpest and strongest absolute v_4_ absorbance followed by *S. clausii* and finally by *L. fusiformis* (Figure 2b). Besides the differences in the vibrational modes, certain bands were evident in biogenic precipitates but absent in synthetic calcite: namely between 950 – 1150 cm^−1^, at 1500 – 1550 cm^−1^, and between 1570 – 1700 cm^−1^. A last band became apparent at 1190 - 1350 cm^−1^, characteristic of amides ^23^, when subtracting the synthetic calcite with the BCC spectra (Supporting Figure 3). Similar spectra have been previously reported on bacterial calcites ^23,43,44^, attributed to overlapping signatures of organics (general organics at 950 – 1150 cm^−1^ and amides at > 1500 cm^−1^) and ACC (v_1_-mode 1060-1080 cm^−1^; a v_3_ asymmetric vibration at 1470 cm^−1^; and O-H bending at 1650 cm^−1^ from structural H_2_O in ACC) (for details see Supporting Information). As such, Mehta and colleagues ^43^ concluded that it is difficult to infer the presence of ACC from the bands at > 900 cm^−1^ alone.

Following the bleaching process used to remove organic matter, BCCs produced almost matching FTIR spectra (Figure 2c). No shift was observed in the carbonate v-modes, yet all three modes experienced an increase in absorbance. This is important as changes in calcite v-mode absorbance, most notably the increase in intensity of the 712 cm^−1^ band, is associated with an increase in crystallinity of the carbonates ^41^. However, the lack of shift in the v_2_ band, and in particular the persistence of the right broad tail, suggests that some ACC was still present, most likely structurally bound to the BCC calcite structure. Conversely, the increase in v_4_ absorption after bleaching suggests that a fraction of ACC, likely the superficially exposed (see TEM results below), was transformed into calcite via dissolution-precipitation in the bleach solution, and/or was removed after rinsing of the bleach. Moreover, the bleaching process leads to removal of bands at higher wavenumbers (bands at 1500 – 1550 cm^−1^ and 1570 – 1700 cm^−1^), previously linked to organics. On the contrary, the bands at 950 – 1150 cm^−1^ and 1190 – 1350 cm^−1^ also corresponding to organics and ACC were conserved after bleaching, with a reduction in absorbance, especially in *L. fusiformis*. Interestingly, the amide-specific band at 1190 – 1350 cm^−1^ in *L. fusiformis* BCC experienced a shift to higher wavenumber after bleaching. Given that we can associate the wavenumber to the bond strength, a higher wavenumber means that a stronger bond exists between the remaining organics and the crystal structure ^45^. It follows that bleaching removed the material more weakly associated with the BCC crystals (e.g. adsorbed organics and ACC), and the subsequent rinsing of the bleach led to dissolution of precursor ACC and recrystallization of calcium carbonate on certain preferential faces of the crystals. This would be responsible for the disappearance of the two humps in the XRD patterns and the changes in Bragg peak intensities. Nevertheless, a fraction of organics (and likely ACC) were not affected by the bleaching process, as a signal of organics remains in the FTIR spectra. Duplicates of the BCCs before and after bleaching show consistent FTIR bands (see Supporting Information and Supporting Figure 4).

Thermogravimetric analysis (TGA) was performed on the bleached BCCs of *L. fusiformis* and synthetic calcite to further disclose the compositional nature of the biominerals (Figure 2e). The calcite control shows a weight loss of 42.3 % starting from 650 °C, close to the theoretical 44 % due to the decarbonation of calcite. The bleached BCC of *L. fusiformis*, showed a continuous and step-wise mass loss at *T* < 100 °C, 150 – 500 °C, 500 – 600 °C and 600 – 950 °C. The initial mass loss is from adsorbed water. The second loss can be from ACC dehydration and/or thermal decomposition of the remaining organic matter. The dehydration of ACC occurs at 100 – 350 °C ^42,46^ and the decomposition of organic matter starts at 250 °C and continues up to 400 or 600 °C for intracrystalline matter ^42,47^. Although there is an overlap between both decompositions, the continuous mass loss after 350 °C distinguishes the presence of organics. Finally, unlike the control, the weight loss of BCC from 650 - 950 °C (33.9 %) does not match the value corresponding to the theoretical mass loss of CaCO_3_ following release of CO_2_, but rather the weight loss from 550 – 950 does (44.1 %). A lower decomposition temperature in CaCO_3_ is associated with lower crystallinity ^48^, which corresponds to the lower crystallinity of BCCs observed with FTIR.

Overall, it becomes apparent that BCCs produced by the two bacteria studied, when formed, are associated with bacterially-derived organic matter and ACC, with distinctive biosignatures in all tests. FTIR confirmed that bleaching removes a non-calcite fraction, seemingly composed of organics and ACC loosely bound to the crystal surfaces, as seen on the SEM micrographs. But FTIR and TGA results show that there is a non-removable organic/ACC fraction that is more permanently associated with the BCC crystals. Overall this produces biomineralized calcite with non-perfect crystal lattice (i.e. lower crystallinity).

### Organics lead to the formation of a distinctive nanostructural Swiss-cheese like structure

To further disclose how the organics were associated with the precipitates, TEM and HRTEM analyses were performed, complemented with HAADF-EDS. Under the TEM we observed that unbleached *L. fusiformis* precipitated crystalline (calcite) and amorphous (ACC) phases as confirmed by selected area electron diffraction (SAED) patterns (Figure 3a, Supporting Figure 5a). Conversely, we could not find individual or aggregated ACC particles in the case of *S. clausii* which only showed single crystal SAED patterns of calcite (Figure 3c). Additionally, SAED patterns with calcite single crystal features were frequently arced, with spots displaying an angular spread of ∼4 to 12° on both BCCs (Figure 3c, Supporting Figure 5a). Angular spreading is a distinctive feature of mesocrystals reflecting a slight misorientation among the different nanocrystals making up a mesocrystal ^49^ and commonly associated with the presence of organic matter in biominerals ^50^. The elemental composition of the unbleached precipitates was confirmed with EDS mapping. Ca, O and C were ubiquitous in all samples. Moreover, it was possible to correlate the presence of organic-related elements such as, N, S and P – the latter two in trace amounts – in the crystals produced by both bacteria (Figure 3b, d, Supporting Figure 5b, Supporting Figure 6).

**Figure 3.**
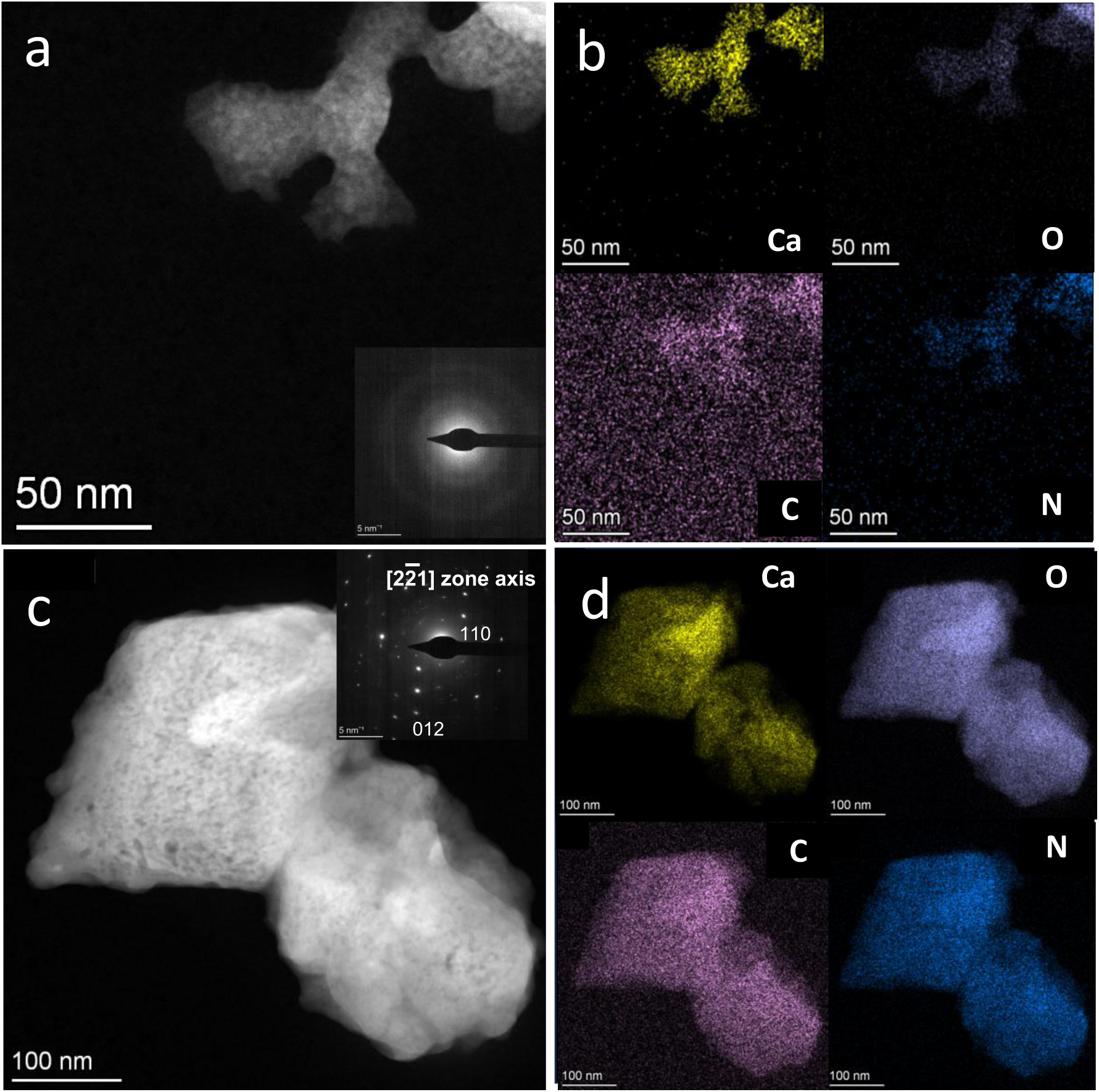
TEM and EDS analysis of bacterial calcium carbonates. (**a**) HAADF image of ACC precipitated by *L. fusiformis*. Inset: SAED pattern showing the amorphous nature of the particle. (**b**) In clockwise order: EDS elemental maps of precipitates in (a) (calcium, oxygen, nitrogen and carbon, respectively). (**c**) HAADF image of calcite crystals precipitated by S. clausii. Note the nanoporous structure of the crystals. Inset: [22̅1] zone axis SAED pattern of the calcite particle on the left with diffraction spots showing an angular spreading of 6°; (**d**) In clockwise order: EDS elemental maps of precipitates in (d) (calcium, oxygen, nitrogen and carbon, respectively).

EDS line-scans from the edge to the interior of the unbleached BCC crystals showed that N was not only present within the crystals but also adsorbed on their surface as the intensity did not drop to zero outside the crystal (Figure **4**a-d). Point scans showed a lower N:Ca ratio in the centre of the crystals than in the rim (Figure **4**e). This implies that the internal areas of the precipitate include a lower relative amount of organics than the rim. HRTEM was used to observe the precipitate rim more in detail. In the case of *S. clausii*, lattice fringe images and corresponding FFT showed that, generally, the outermost layer of the bacterial calcite crystals was amorphous, confirming previous observations ^23^ (Figure **4**f-h, more examples in Supporting Figure 7). Moreover, here we also observed amorphous areas, interwoven within the crystalline lattice (Figure **4**f). This occurred in a region of intermediate contrast and a few nanometers in thickness at the edge of the crystal, which was N-enriched according to EDS analysis (N:Ca ratio of 0.53 Figure **4**e). The latter points to occlusion of organics and/or amorphous calcium carbonate, consistent with the relatively high Ca content (Figure **4**e).

**Figure 4.**
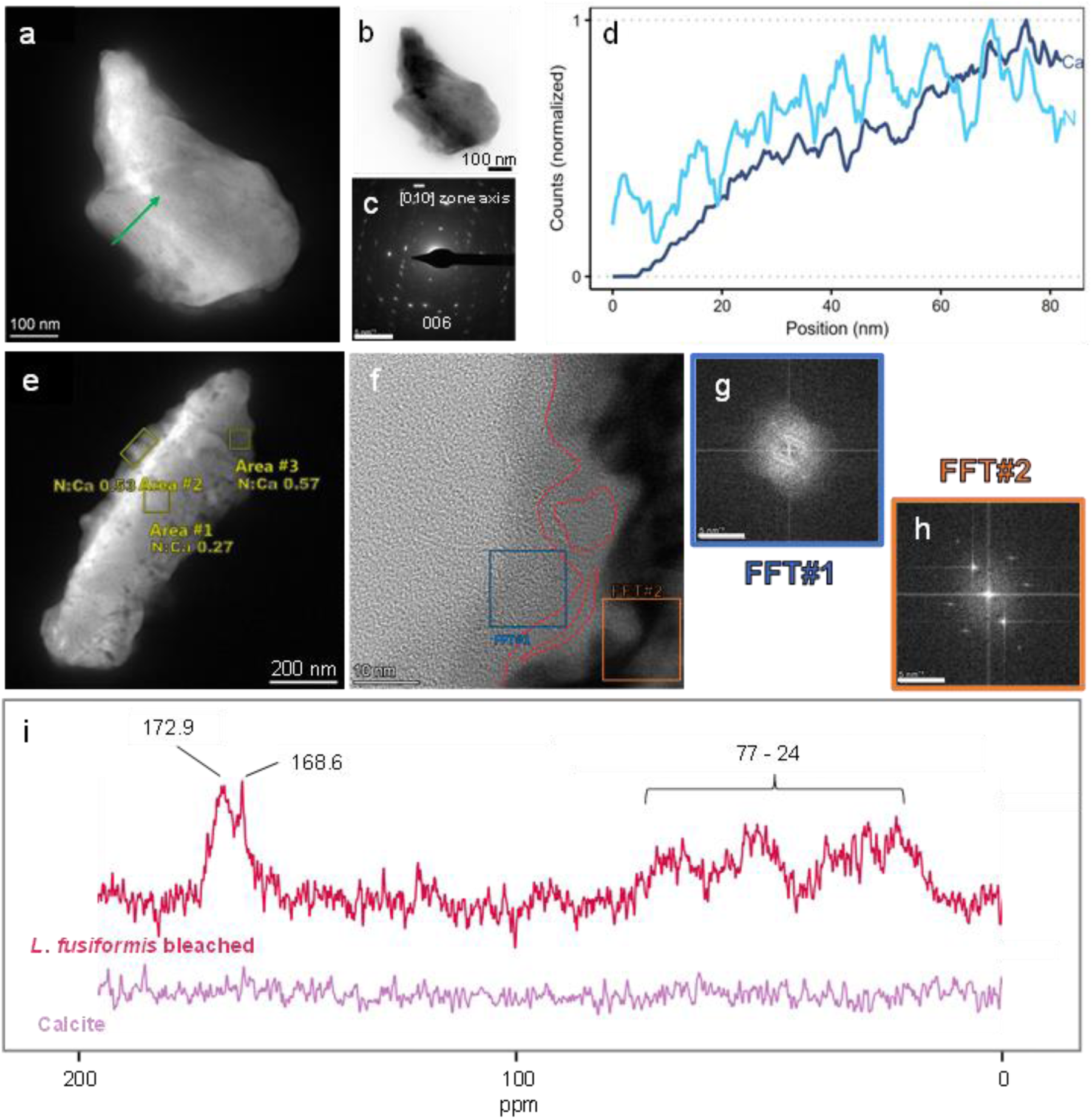
HRTEM, HADDF-EDS and ss-NMR analysis of bacterial calcium carbonates. **(a)** HAADF image of calcite crystal precipitated by *L. fusiformis*. The porous -Swiss-cheese like-structure of the calcite crystal is visible and the location of the EDS line scan is indicated (green arrow). **(b)** Brightfield TEM imagae of the crystal in (a). **(c)** [01̅0] zone axis SAED pattern of the crystal shown in (a) and (b). Note the marked arcing of diffraction spots (showing an angular spreading of ∼9°). **(d)** Line scan in (a) showing that N concentration (light blue line) is uncoupled (after 40 nm) to that of Ca (dark blue line), and the increase in N can be traced to the pores on the calcite crystal (corresponding to the darker areas in (a)). **(e)** *S. clausii* calcium carbonate structure showing different N/Ca ratios in different areas. **(f)** HRTEM image of the edge of the precipitate in (e) (Area #2) with FFT of the blue squared area **(g)** demonstrating it is an amorphous external layer, and the FFT of the orange squared area **(h)** showing it is a crystallized (calcite) internal zone. The red outline demarcates the end of the lattice fringes, showing amorphous areas trapped within the crystalline zone at the rim of the precipitate. **(i)** CP–MAS ^13^C NMR spectra of bleached *L. fusiformis* calcite (light red) and synthetic calcite (purple). The peak at 172 ppm is associated with calcite while the remaining peaks are due to the presence of amides. Referenced to the signal of DSS (δ = 0 ppm).

Furthermore, an uncoupling of N and Ca was observed on the line-scans in the inner region of the crystals (> 40 nm from the edge), i.e. when Ca increased, N decreased (Figure ***4***d), albeit with a slight phase-shift. Upon close inspection of HAADF images, it was clear that the crystals did not show a homogenous surface (Figure ***4***a). Rather, a Swiss cheese-like structure could be observed, with elongated generally parallel low Z-contrast pore-like areas present within the bacterial calcite single crystals (more examples in Supporting Figure 8). A direct association of these low Z-contrast nanoporous-looking areas with an increase in N content was systematically observed. It could be argued that this porous structure is the result of beam damage. However, this is ruled out here because the HAADF images were collected before EDS analysis and the porous structure was already present. Pioneering work by Towe and Thompson ^51^, and many other subsequent studies ^52^ have also shown that beam damage does not produce such features, but organics do. Moreover, this structure was also observed in many other BCCs during brightfield TEM analysis and subsequent SAED analysis showed no CaO diffraction spots/rings which are standard features of beam damage in calcite ^53^. Note also that we have not observed such Swiss-cheese structure in any studied synthetic calcite crystal. Although a Swiss-cheese-like structure has never been reported in the case of BCC, similar features have been observed in other calcium carbonate biominerals, such as mollusc shells ^52,54^, and their biomimetic counterparts ^55^. The pore-like areas have been usually identified as intracrystalline organic matter and recently with ACC (see Supporting Information).

CP-MAS ^13^C NMR (ss-NMR) and HRXRD were used to confirm the nature of those observed here, utilizing bleached precipitates. The CP-MAS ^13^C NMR spectra of bleached BCC primarily consist of broad peaks, with a notable exception at 168 ppm attributed to calcite (Figure ***4***i), as confirmed by the MAS ^13^C of synthetic calcite (Supporting Figure 9). The broad peaks are indicative of amorphous content yet no well-defined peak was observed closely associated downwards of calcite (167 ppm), where ACC is typically present ^56^. The remaining option for amorphous content must then be organic in nature. Many broad peaks are present in the 20 - 80 ppm range and a peak at 172 ppm suggests the presence of carbonyl groups. These shifts are linked to amino acids which have peaks at 20-60 ppm (CH), and 170-180 ppm (carbonyl) ^57^. Overall, the spectra closely resembles that of eggshells ^58^ and protein-including calcite biominerals ^59^. The CP-MAS ^13^C of the synthetic calcite does not show any bands, as expected (Figure ***4***i; see also Supporting Figure 9). This means that the calcite peak present in the *L. fusiformis* BCC is associated with hydrogen atoms, only available from either organics or water ^56.^ Because the presence of water, such as in monohydrocalcite ^56,^ should broaden and shift to higher ppm values the calcite carbonyl bond, we conclude that the hydrogen is from amine or carboxyl groups from the organics. Lastly, the broadness of the peaks further suggests that the structure is internal ^56^. This suggests that the observed Swiss cheese-like structure must be present through the whole crystal and not superficially as seemingly observed on TEM.

To further disclose whether these regions are internal, and related to the organic signatures ubiquitously observed, we performed HRXRD analyses at the CRYSTAL beamline of the Soleil synchrotron (Paris, France). HRXRD analysis showed that the 104 and 006 Bragg peaks of bleached BCC produced by *L. fusiformis* were left-shifted (i.e., to higher *d*-spacings) as compared with geogenic calcite (optical quality Iceland spar) (Figure 5a, b). The shifting was more marked for the 00*l* reflections, implying anisotropic lattice distortion. Rietveld full profile fitting analysis ^27^ yielded the following refined cell parameters for BCC calcite: *a* = 4.98900(4) Å; *c* = 17.08345(5) Å; cell volume = 368.24(4) Å^3^, demonstrating a clear anisotropic lattice distortion as compared with reference geogenic calcite (*a* = 4.98970(2) Å; *c* = 17.05900(4) Å; cell volume = 367.82(2) Å^3^). Maximum lattice strain was observed along the [001] direction (Δ*c*/*c*=0.0014). This lattice strain fluctuation is within the range reported for biogenic (e.g., mollusc shell) and biomimetic calcite ^60–63^. HRXRD analysis during *in situ* isochronous heating (annealing) from room *T* up to 400 °C at 50 °C intervals (see Materials and Methods), showed little changes in Δ*a*/*a* and a significant Δ*c*/*c* reduction at *T* > 200 °C (Figure 5c). Reanalysis of the BCC after annealing at maximum *T* and subsequent cooling to room *T* showed permanent lattice contraction and peak broadening (Figure 5). Pokroy et al. ^60,61^ observed exactly the same features following similar (isochronous annealing) HRXRD analysis of calcite in mollusc shells, demonstrating intracrystalline occlusion of organics (as solutes) in such biominerals. The preferential expansion along the *c*-axis was associated with interaction/replacement of carbonate groups placed along (00.1) planes of calcite by acidic proteins, although interaction with other polar planes such as (01.2) should also be favoured ^60,63^. The authors also stated that thermal decomposition of occluded organics at *T* > 200 °C results in strain relaxation and reduction in Δ*c*/*c*, with a parallel increase in peak broadening due to reduction in crystallite size and increase in microstrain fluctuations, as we have observed here. Interestingly, we also observed an increase in Δ*c*/*c* of BCC between 100 °C and 200 °C. The latter, also observed in other calcite biominerals such as sea-urchin spicules, has been associated with the presence of ACC and its thermal-induced transformation into calcite, which leads to expansion ^64^. Altogether, our HRXRD results confirm that intracrystalline organic occlusions also occur in bacterial calcite, with very similar anisotropic lattice distortion as in other calcite biominerals such as mollusc shells. Our HRXRD results also suggest that intracrystalline ACC is present in the BCC structures, as supported by TEM and ss-NMR results.

**Figure 5.**
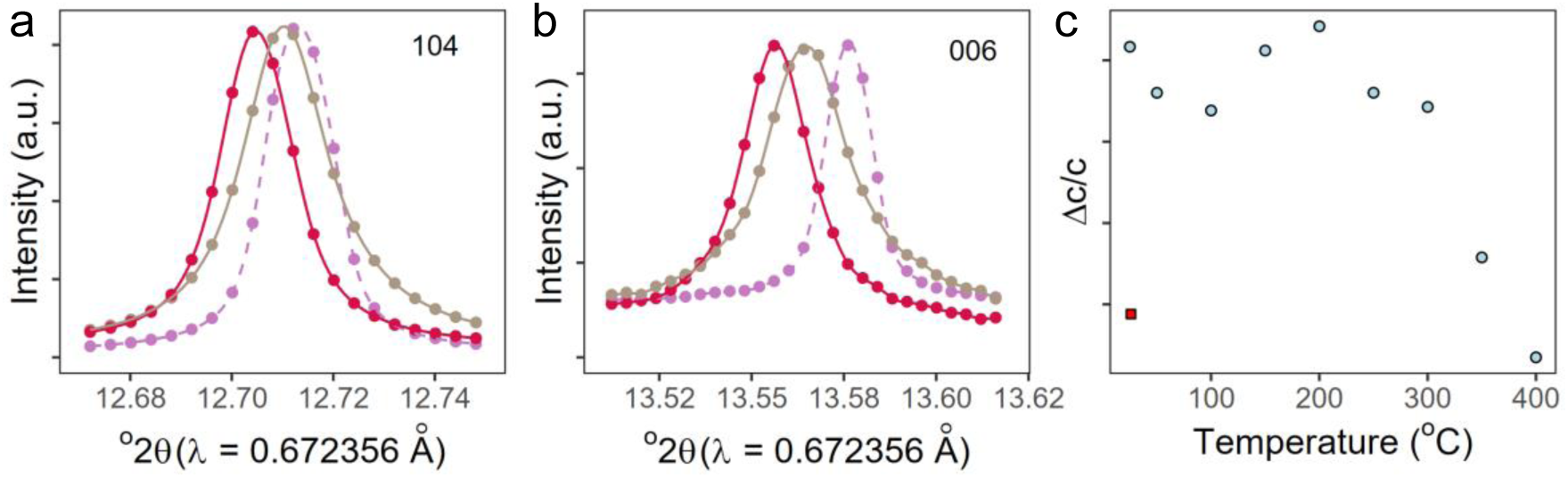
Synchrotron HRXRD analysis of *L. fusiformis* calcite. **(a)** 104 Bragg peaks and **(b)** 006 Bragg peaks of bacterial calcite before (red) and after annealing at 400 °C and cooling to room *T* (light brown). The dashed (purple) profiles show the reference Iceland spar calcite; **(c)** *T*-dependent variation of lattice distortion (Δ*c*/*c*) along the *c-*axis as compared with reference geogenic calcite. The red squared symbol marks the residual lattice distortion after annealing at 400 °C and cooling to room *T*.

### Role of crystalline 2D organic films in BCC formation

During TEM imaging of *L. fusiformis* biogenic precipitates we unexpectedly observed layered structures containing areas of amorphous and crystalline carbonates (Figure 6a). HRTEM imaging showed that these film-like structures were crystalline displaying lattice fringes oriented in different directions. SAED analysis of the film-like structures revealed diffraction spots with *d*-spacings 3.7 – 4.5 nm aligned in rows with different orientations corresponding to the multiple orientations of the lattice fringes visualized in the HRTEM image (Figure 6a,b yellow and green lines), along with diffraction spots corresponding to (at least two non-oriented) calcite crystals (Figure 6c). The *d*-spacings of the films is at the scale of those measured in protein crystalline structures ^65^ but full characterization of these structures to confirm if they were made up of proteins was not possible. HRTEM images showed the carbonates closely associated with these 2D crystalline films. The crystalline films were less visible on higher density patches indicating that they were placed at the “bottom” of calcite. However, as shown in Supporting Figure 10a, we could observe in other examples a close relationship between the film and an overlying nanogranular structure, with a SAED pattern corresponding to a calcite single crystal (Supporting Figure 10b). These results suggest that the formation of oriented calcite after particle attachment of ACC is templated by the film underneath, in a similar fashion as that reported on S-layers ^66^ or carbonic anhydrase films ^67^ (see Supporting Information).

**Figure 6.**
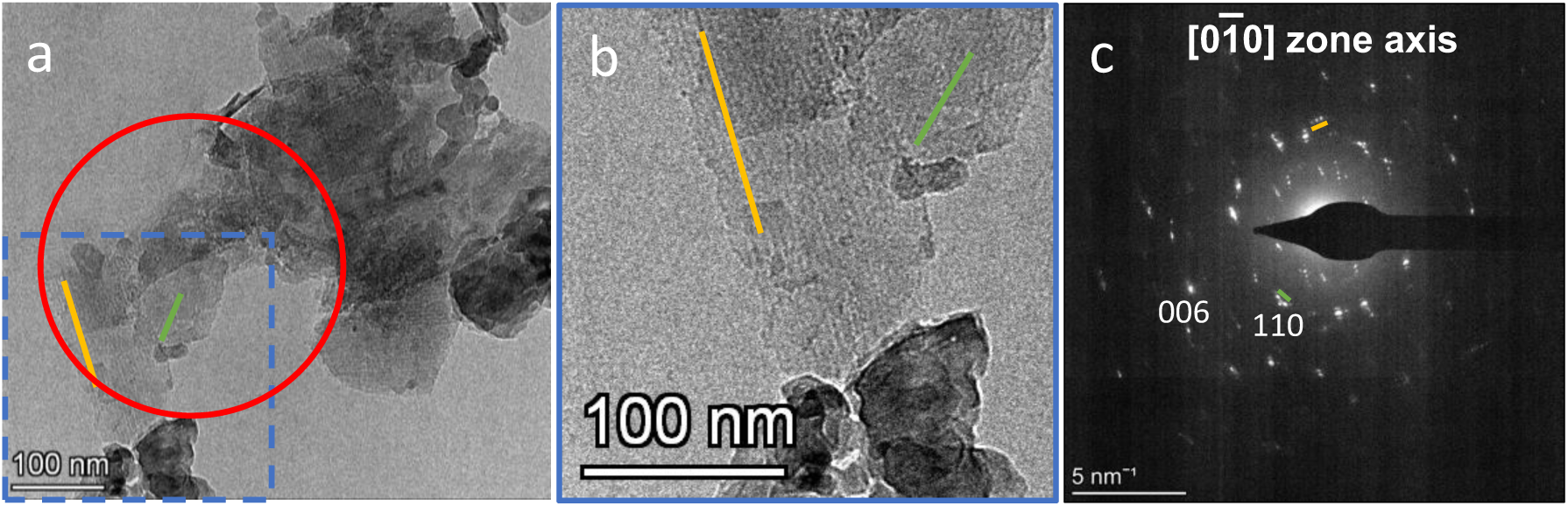
HRTEM analysis of *L. fusiformis* calcite formed on an organic crystalline 2D template. **(a)** A polycrystal showing different amorphous and crystalline phases, alongside crystalline 2D film showing lattice fringes. **(b)** Zoom-in of the blue squared area in (a) showing lattice fringes in the 2D films. The yellow and green lines show the direction of the lattice fringes. **(c)** SAED pattern of the red circled area in (a) showing different diffraction spots corresponding to calcite ([01̅0] zone axis, calculated using the most intense diffraction spots: note that additional orientations are present) overprinted with diffraction spots of the (organic) films (the green and yellow lines mark the three aligned spots corresponding to the films in (b) with lattice fringes oriented along the green and yellow lines).

## Discussion

FESEM showed that the bacteria were closely associated with the BCC crystals and that a nanogranular texture was present, suggesting the formation through an ACC precursor. XRD indicated that all carbonates were calcite, while amorphous and organic signatures were detected with FTIR and TGA in both bleached and unbleached samples. TEM analysis showed the presence of both crystalline and amorphous calcium carbonate, the latter being more predominant in *L. fusiformis*, in agreement with FTIR spectra. In the individual crystals, amorphous areas were present at the rim of the BCCs, likely the growth front of the crystal, in both bacteria. TEM-EDS mapping unveiled the presence of organic-related elements in the bacterial crystals, including N, (and S and P to a lesser extent) associated with both crystalline and amorphous regions of calcium carbonate precipitates. Importantly, we observed low Z-contrast pore areas that resembled a Swiss cheese-like structure thoroughly reported in calcifying metazoans ^26,54^, and a strong correlation with N was detected in the low Z-contrast regions. CP-MAS C^13^ NMR pointed at the presence of amino acids in a strikingly similar way as in other protein-including calcite biominerals ^59^. TEM-EDS and ss-NMR results helped clarify that the observed FTIR bands correspond to amide bonds (Amide I band (1600 – 1700 cm^−1^); Amide II: 1540 cm^−1^ likely attributed to a C=N bond ^68^ and C-N-C=O bonds at 1499 cm^−1 69^; Amide III band 1250 – 1350 cm^−1 69^). The overall strong indication of amides points to peptides having a role in the biomineralization process, although no particular macromolecule was determined.

The detection of organic signatures by FTIR and TGA after bleaching suggests that an organic fraction remained bound to the BCCs. Three possibilities exists for this explanation: (i) the bleaching is not sufficient to remove all organics from the precipitate surface, (ii) the organics can also be structurally bound and protected from the bleaching by being occluded in the precipitates, or (iii) both. The presence of organic occlusions in BCCs is as of yet unreported and could have important practical implications while changing our understanding of how biomineralization in bacteria works (see below). While ss-NMR supports the second hypothesis, HRXRD unambiguously shows that this fraction was (in part), intracrystalline organic occlusions. HRXRD shows that the amino acids/peptides are intracrystalline, as there was a marked lattice strain (expansion) along the *c*-axis, and a lattice contraction was observed along the *c*-axis direction after annealing (up to 400 °C). Similar behaviour has been observed using HRXRD in the case of calcite biominerals produced by eukaryotes and their biomimetic counterparts ^60–62^. Note that the nanometer-sized porous areas filled with N-rich organics forming the Swiss-cheese structure are, however, too large to account for the lattice strain fluctuations determined by HRXRD. Apparently, organics are occluded in BCCs at two scales: the atomic scale (as solutes in the calcite crystal lattice) resulting in the observed lattice strain, and the mesoscale (as nanometer-sized clusters) as observed here with TEM. This latter nanometer size occlusion could be responsible for the angular spreading observed in SAED patterns and become occluded as the crystal grows at the rim. This is consistent with the amorphous phases occluded inside the crystal lattice observed in HRTEM analysis. Similar organic occlusion occurs in several calcium carbonate biominerals produced by eukaryotes ^52^. Note that foreign nanoparticle occlusion results in peak broadening due to microstrain contributions associated with the creation of intracrystalline boundaries acting as a source of inhomogeneous deformation fields but without necessarily contributing to macrostrain, i.e., lattice distortion ^61^. These results are consistent with the ss-NMR results as the broad peaks of the CP–MAS ^13^C NMR spectra point to intracrystalline presence of the amino signatures. Finally, the confirmed role of organics in the crystallization process, explain the preferential growth of BCC calcite along the *c*-axis, with overdeveloped {110} (pseudo)faces, which are typically formed during calcite growth following adsorption of organics on acute steps ^70^. Besides the organic signatures, we also report strong indications of ACC presence throughout the whole study, even after bleaching. If the organics remain occluded, then this suggests that ACC is also occluded, and likely stabilized by the organics as observed elsewhere^37^. However, further analysis is needed to unambiguously prove this such as that presented by De Frutos et al. ^26^.

The evidence presented here supports the mechanism of precipitation originally alluded to by Enyedi et al. ^23^, where ACC is stabilized by EPS, and calcite forms through PA. However, we clarify several missing aspects after ACC stabilization. Firstly, we propose that BCC grows by aggregation of protein-stabilized ACC, rather than the more broadly assigned EPS. This is consistent with the intracrystalline occlusion in BCC calcite crystals of both peptides and ACC. Moreover, we observed that larger crystalline 2D films structures, not only stabilize, but act as a template for crystal formation. This film had *d-*spacings in the range of those observed in crystalline protein films, although the molecular features of the films could not be fully characterized. The stabilized ACC then undergoes an amorphous-to-crystalline conversion, at which point the intracrystalline occlusion of a fraction of the organics associated with ACC occurs. This closely resembles the current model of biomineralization of calcite biominerals in eukaryotes ^26^. Our proposition is in contrast to what was previously suggested for bacteria, where organics were assumed to be completely ejected ^23^. Biomineralization through the eukaryotic model involving an ACC precursor commonly produces a nanogranular structure and results in a Swiss cheese-like structure, as observed here. Even more remarkable, de Frutos and colleagues proposed a model for calcite biomineralization in animals wherein pellicle-like regions of concentrated organics are formed ^26^. These are starkly similar to the orderly fashion in which the pore-like N-rich structures were observed in our bacteria biominerals. Moreover, the transformation of ACC nanoparticles into calcite occurs in crystallographic register with the core calcite, which explains the single crystal features of the SAED patterns. Still, some miss-orientation occurs, resulting in the observed angular spreading and the mesocrystal features described above, which are also observed in calcite biominerals produced by eukaryotes, as it is the case of sea-urchin spicules ^71^. This could point towards a shared biomineralization process, but future research needs to explore such hypothesis.

On top of other calcium carbonate biominerals, similar structures as those observed for the BCCs in this work form in biomimetic *in vitro* models ^55^. This might not be surprising given that many processes proposed for the formation of calcite biominerals have been shown to be thermodynamically favourable (i.e., down-hill energy landscape) ^36^. Yet, critically, in these models the carbonate crystals do not display the highly organized organic-inorganic nanostructure observed in biominerals, in general, and in our BCCs, in particular. Thus, the organization of the BCCs mesocrystals and the hierarchy in organic occlusions observed in this work could contradict the common idea that calcium carbonate biomineral precipitation by microorganisms - in contrast CaCO_3_ biominerals produced by eukaryotes - is a spontaneous and uncontrolled phenomenon ^8^. Furthermore, other authors are already scrutinizing the idea that calcium carbonate biomineralization in prokaryotes is functionless ^72^. Some recent work points to real adaptive benefits for MICP. For example, Botusharova et al. ^12^ showed that bacterial spores could survive entombment in carbonate precipitates, being autoclaved and then reactivated. More recently, it was shown that the precipitation of calcium carbonate had a central role in the effective formation of biofilm in pathogenic bacteria with genetic inhibition of carbonate biomineralization leading to a reduced pathogenicity and biofilm formation ^10^. On the other hand, control over the precipitation process could provide protection to the bacteria from unfavourable conditions by controlling the rate and location on the cell where precipitates form, for example to prevent unwanted entombment ^66^, or reduce damage from toxic metals by removing these from the surrounding environment through (co)precipitation in carbonates ^73^. Although the work remains limited, investing energy in producing organics that will create a controlled process of biomineralization might contribute to the inclusive fitness of the organism.

Aside from the evolutionary standpoint, there is technical applicability to these results. The observations of intracrystalline organics and possibly also ACC, implies that there might be improved mechanical properties for BCCs. Intracrystalline organic matter that occurs in other calcium carbonate biominerals such as mollusk shells, lead to improved mechanical properties (e.g. increased toughness) ^52^. Moreover, stabilized ACC as a precursor prior to transitioning to calcite has been shown in different biomineral composites to lead to improved structural properties ^74^. This observations can have implications in the technical applications of microbial biominerals, such as in bioengineering applications of MICP, like self-healing cements or the consolidation of stone built heritage. Moreover, identification of these biosignatures could pave the way for determining biogenicity of carbonate samples in the geologic record, which is not always straight forward. As such, understanding how these mechanisms work and are controlled can open up the door for novel biomaterial applications with purposely designed structure-function.

## Supporting information

All supplementary data

## Supporting information

Additional experimental details and results including Supporting Figures 1 – 10 and Supporting Table 1.

## Acknowledgments

We thank the personnel of the Centro de Instrumentación Científica (CIC) of the University of Granada for helping with FESEM and TEM analyses and the Inorganic Chemistry department for access to their FTIR. Additional thanks go to the personnel of the Department of Mineralogy and Petrology of the University of Granada for their help with the instrumentations, including Teodora Ilić, Sarah Bonilla Correa, Aurélie Pace and Miguel Burgos Ruiz. Special thanks go to Kerstin Elert for her help with the SEM imaging and Fadwa Jroundi Mesbahi for allowing to grow our bacterial cultures at her facilities. HRXRD experiments were performed at CRYSTAL beamline at SOLEIL Synchrotron with the help of this beamline staff (E. Elkaim). We also thank Julia Atienzar, Aurelie Pace, and Nuria Pujol Solà for their help during HRXRD analyses at SOLEIL synchrotron and Alejandro Rodriguez-Navarro for his help during Rietveld analysis of HRXRD results.

## Author Contributions

FGG contributed Conceptualization, Methodology, Investigation, Visualization, Supervision, Writing (original draft), Writing (review and editing)

MP contributed Conceptualization, Investigation, Visualization, Writing (original draft), Writing (review and editing)

NB contributed Conceptualization, Supervision, Writing (review and editing), Funding acquisition

NDB contributed Conceptualization, Supervision, Writing (review and editing), Funding acquisition

CR-N contributed Conceptualization, Methodology, Investigation, Visualization, Supervision, Writing (original draft), Writing (review and editing), Funding acquisition

