## Supplementary material for "Ultrastructural features of bacterial calcite unveil ubiquitous organics occlusion and clues on biomineralization": All supplementary data

This PDF file includes:

Supporting Information

Supporting Figures 1 to 10

Supporting Tables 1

SI References

### Supporting Information

#### Culture and precipitation media, and precipitate extraction

The preparation of media in glassware could lead to silica dissolution and contamination of the precipitates. We note, however, that none of the observed bands correspond to $SiO_2$. To test this, liquid B4 was prepared by autoclaving in glassware, which could release silica into the system, or through filter sterilization. Then either media was inoculated in a glass Erlenmeyer or a plastic Falcon tube. The crystals were then analyzed using FTIR. Media preparation had no effect on the bands produced (Supporting Figure 1). This is further supported by the lack of silica identification with Raman (characteristic at 520 $cm^{-1}$ and 480 $cm^{-1}$ for crystalline and amorphous, respectively).

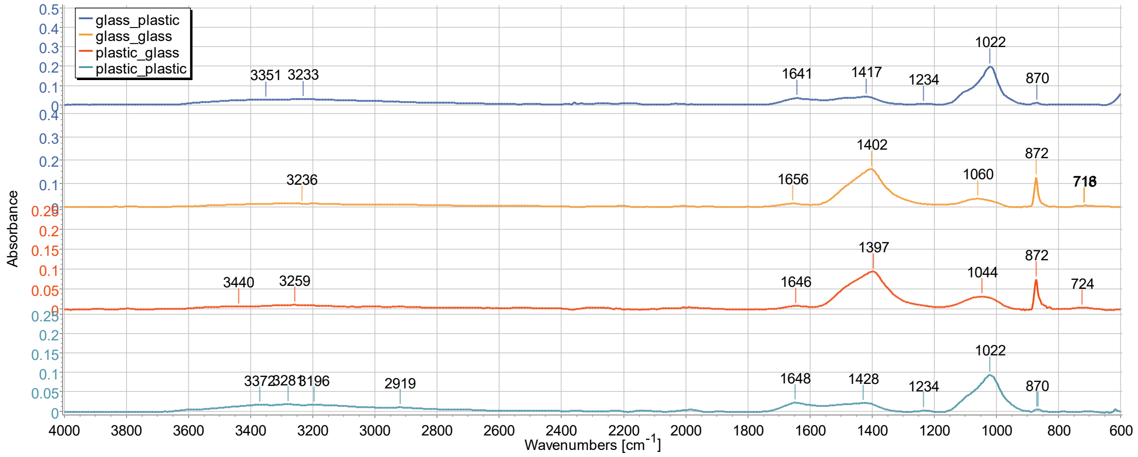


**Supporting Figure 1. FTIR of *L. fusiformis* crystals from media prepared in glassware and autoclaved (glass), or in plastic and filter sterilized (plastic).** The container type in which the inoculation was performed is also indicated (_glass or _plastic).

**Bleaching of precipitates**

To ensure that 2h of bleaching was robust to remove all organics, aliquots of crystals were incubated for 10 min, 30 min, 2 h, 4 h and 8 h. The resulting crystals were checked with FTIR. Overall it was clear that after 10 min, the FTIR spectra were showing that all bands that were not bound to the crystal structure were already removed (Supporting Figure 2).

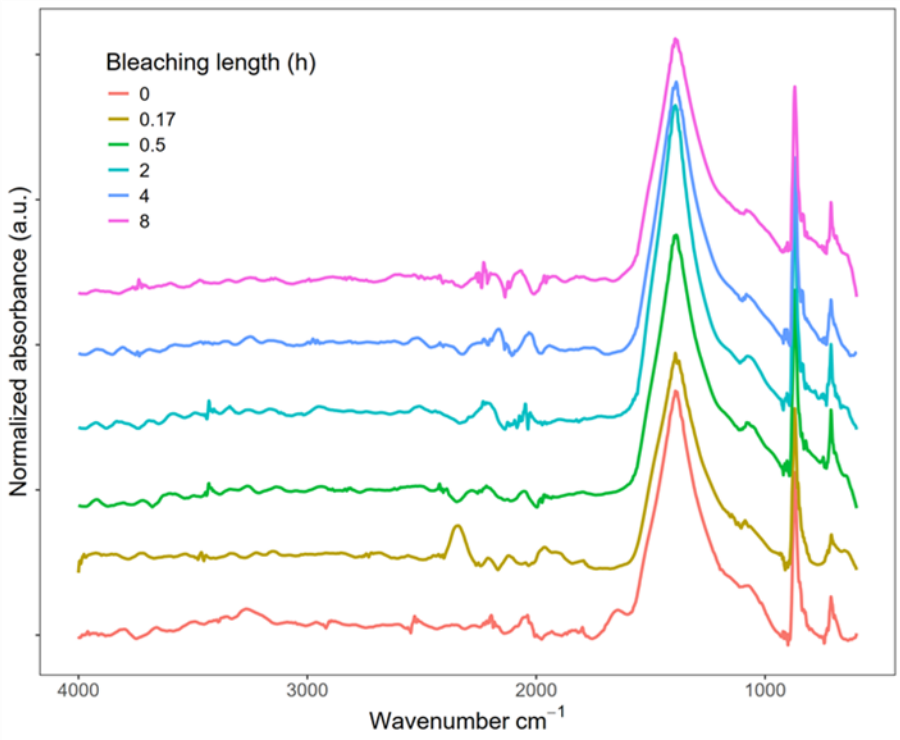


**Supporting Figure 2. FTIR spectra of *L. fusiformis* calcite through different bleaching times from unbleached (0) until 8 h.** The bands at 1500 – 1700 cm$^{-1}$ are removed already after 10 min (0.17 h) of bleaching and the spectra remain the same after that.

**Additional XRD results**

Overdevelopment of the {11.0} form results in preferred orientation during powder XRD analysis, leading to a relative increase in the intensity of the 110 Bragg peak of calcite (as compared with the most intense 104 Bragg peak) as shown in Table S1 for the case of BCCs. The overdevelopment of the {11.0} form is characteristic of calcite biominerals, as well as biomimetic calcite formed in the presence of different organic additives. Previous research on other biomineralized calcium carbonates has shown that, despite the (11.0) face's higher surface energy (as compared with (10.4) faces), this change in morphology is due to the preferential adsorption of a range of organic molecules on such faces. Note, however, that Teng and Dove (5) indicate that such faces actually are pseudo-faces composed by (01.2) and (11.0) faces. Note also that (01.2) faces are polar and expose alternating layers of calcium and carbonate ions. This also occurs with (00.1) faces, where alternating layers of calcium and carbonate ions are stacked along the *c*-axis. Such polar faces are optimal for the interaction with the functional groups of a range of (macro)molecules such as (acidic) proteins, DNA, amino acids, and polysaccharides, among others (6–8). For example, carboxylate additives led to the precipitation of spindle shaped crystals displaying overdeveloped (11.0) (pseudo)faces (9). Zhu et al. (10) showed how acrylic acid easily replaces water on the (11.0) plane allowing for stronger organic-crystal interactions than with the (10.4) planes thus resulting in the preferential growth of calcite along the *c*-axis with overexpressed (11.0) faces. Davila-Hernandez et al. (11) designed a protein template that helps assemble calcite with preferred crystallographic orientation and displaying non-natural (11.0) or (20.2) faces. As such, the observed relative increase in the intensity of the 110 Bragg peak, strongly suggests the involvement of organics in the process of microbial crystallization of calcite. Furthermore, crystals displaying overdeveloped (11.0) (pseudo)faces have been connected to slow and controlled ACC-to-calcite transitions such as at high temperature (12), or in biomimetic synthesis of calcium carbonate films (13).

Table S1 - Relative intensities (Rel. Int.) of the main *hkl* Bragg peaks and their °2θ position (Pos.) of each XRD pattern (Cu Kα radiation).

| Synthetic calcite | | | *L. fusiformis* precipitates | | | *S. clausii* precipitates | | |
|---|---|---|---|---|---|---|---|---|
| Pos. [°2θ] | *hkl* | Rel. Int. [%] | Pos. [°2θ] | *hkl* | Rel. Int. [%] | Pos. [°2θ] | *hkl* | Rel. Int. [%] |
| 29.4 | 104 | 100.0 | 29.4 | 104 | 100.0 | 29.4 | 104 | 100.0 |
| 36.1 | 110 | 3.7 | 35.9 | 110 | 33.3 | 36.0 | 110 | 59.2 |
| 39.4 | 113 | 4.3 | 39.4 | 113 | 27.1 | 39.4 | 113 | 26.3 |
| | | | *L. fusiformis* bleached precipitates | | | *S. clausii* bleached precipitates | | |
| | | | 29.4 | 104 | 100.0 | 29.4 | 104 | 88.2 |
| | | | 35.9 | 110 | 37.0 | 36.0 | 110 | 100 |
| | | | 39.4 | 113 | 27.7 | 39.4 | 113 | 31.0 |

**Additional FTIR results**

A detailed description of the identified FTIR bands in Figure 2B is outlined here. The first prominent band had a maximum at 1064 $cm^{-1}$. This range corresponds to C-O bonds (1060 $cm^{-1}$) (14, 15), P=O bonds (1080 $cm^{-1}$) (16) and O-H bonds (1082 $cm^{-1}$) (17), found in many organic molecules including polysaccharides and proteins. At the same time, the $v_1$ symmetric stretching of ACC can be identified at 1060-1080 $cm^{-1}$ making it very hard to accurately identify this range using solely FTIR (14, 18). The second band, between 1500 – 1550 $cm^{-1}$ and present only for *L. fusiformis*, strongly overlaps with the Amide II band (1510 – 1580 $cm^{-1}$). Within this band, two maxima are present: one at 1540 $cm^{-1}$ likely attributed to a C=N bond (19) and at 1499 $cm^{-1}$ from C-N-C=O bonds (17), both Amide II groups. It's worth noting that ACC also exhibits a $v_3$ asymmetric vibration band at 1470 $cm^{-1}$, which may influence this region (14, 18). Finally, the 1570 – 1700 $cm^{-1}$ band present in both bacterial precipitates covers the whole range of the Amide I band (1600 – 1700 $cm^{-1}$), and is mainly associated with the C=O stretching found at 1630 $cm^{-1}$, precisely where the maximum lies (18–20). Once again, we note that ACC has a band at 1650 $cm^{-1}$ corresponding to O-H bending (i.e., structural $H_2O$ in ACC) (14).

By subtracting the bleached and unbleached FTIR spectra, we observed that a band at 1190 - 1350 $cm^{-1}$ becomes apparent, which was not obvious originally (Supporting Figure 2).

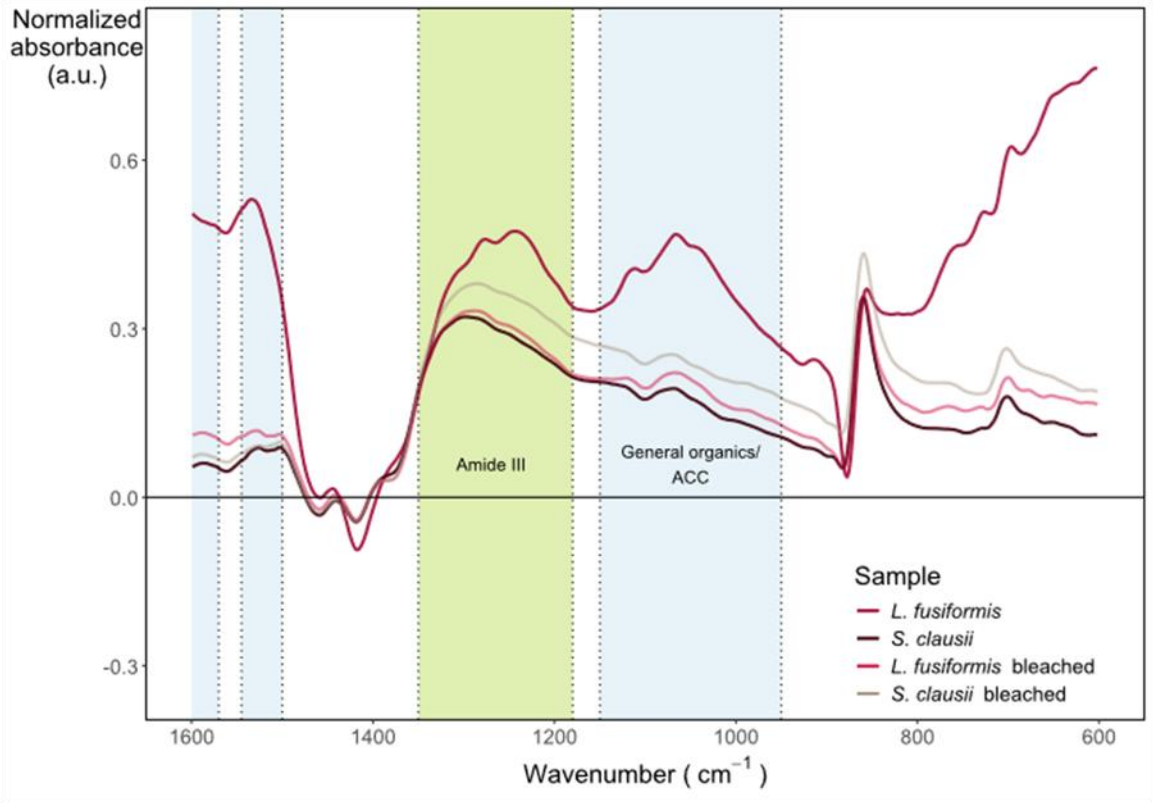


**Supporting Figure 3. Subtraction of FTIR spectra of both biogenic crystals before (red and green) and after (light red and green) bleaching with the FTIR spectrum of synthetic calcite.**

FTIR analysis was performed in duplicate, using newly produced BCC, to disclose whether there could be any variability in the spectral features. Supporting Figure 4 shows the FTIR of the duplicates, both bleached and unbleached. The precipitates showed the same bands, confirming the presence of a mix of organics (strong amide presence) and both calcite and amorphous phases. Subtraction of the spectra produced similar results as well, with the amide bands observable (not shown). However, a stark difference in the absorbance and sharpness for bands at 930 – 1700 $cm^{-1}$ of *L. fusiformis* precipitates was observed*.* This implies that on the second run, *L. fusiformis* produced a lower amount of organic by-products. Given the variability of bacterial kinetics this is not surprising. Nevertheless, we note that the bleached precipitates of the duplicate run show very similar FTIR spectra as those of the original ones, which demonstrates that the compositional/structural characteristics of the produced precipitates are very similar and reproducible.

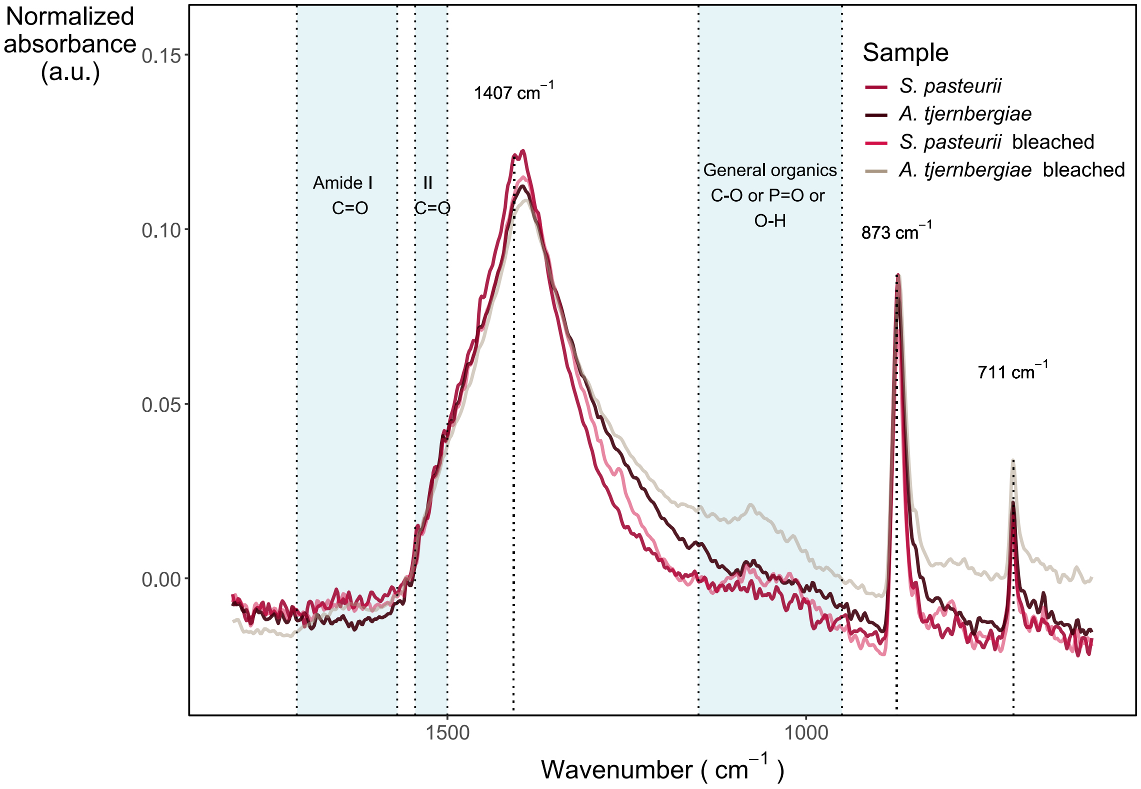


**Supporting Figure 4. FTIR spectra of duplicate *L. fusiformis* and *S. clausii* precipitation experiments.** Spectra show calcite before (red and brown, respectively) and after (light red and light brown, respectively) bleaching. The FTIR spectra of *L. fusiformis* before bleaching fitted more closely to that of *S. clausii* compared to the precipitates obtained in the first precipitation run.

**Additional TEM-EDS result**

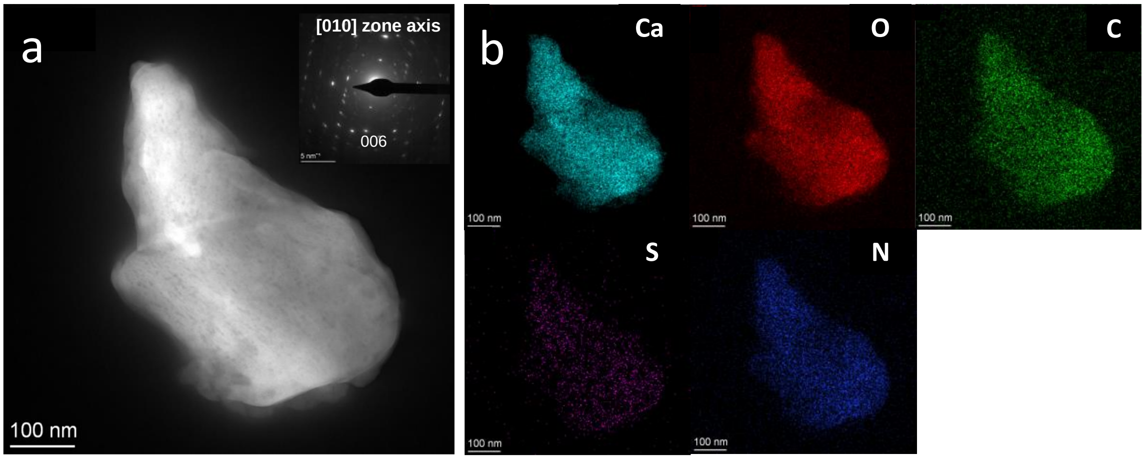


**Supporting Figure 5. TEM and EDS analysis of bacterial calcium carbonates. (a)** HAADF image of calcite crystal precipitated by *L. fusiformis*. The porous structure of a calcite crystal is visible. Inset: [010] zone axis SAED pattern of the crystal. Note the marked arcing of diffraction spot (showing an angular spreading of ~9°). **(b)** EDS elemental maps of precipitates in (a) (calcium, oxygen, carbon, nitrogen and sulfur, respectively).

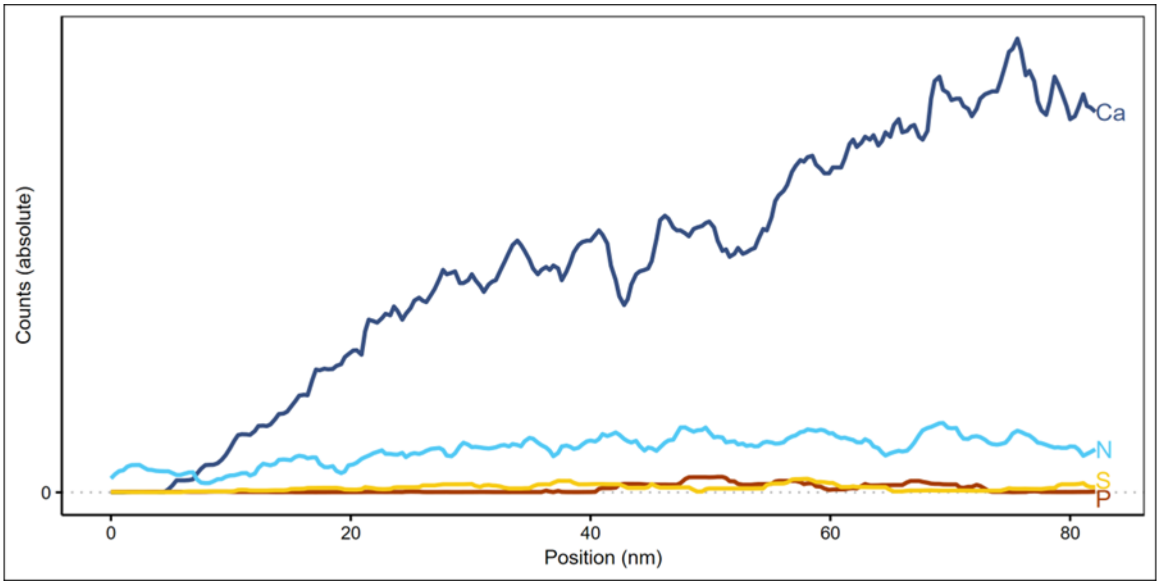


**Supporting Figure 6. Line scan corresponding to the green arrow depicted in Figure 3a.** showing that Ca, N, S and P are present on the crystalline precipitates of

*L. fusiformis.* N was the most dominant organic element. Note these are absolute counts vs normalized counts shown in Figure 4 d.

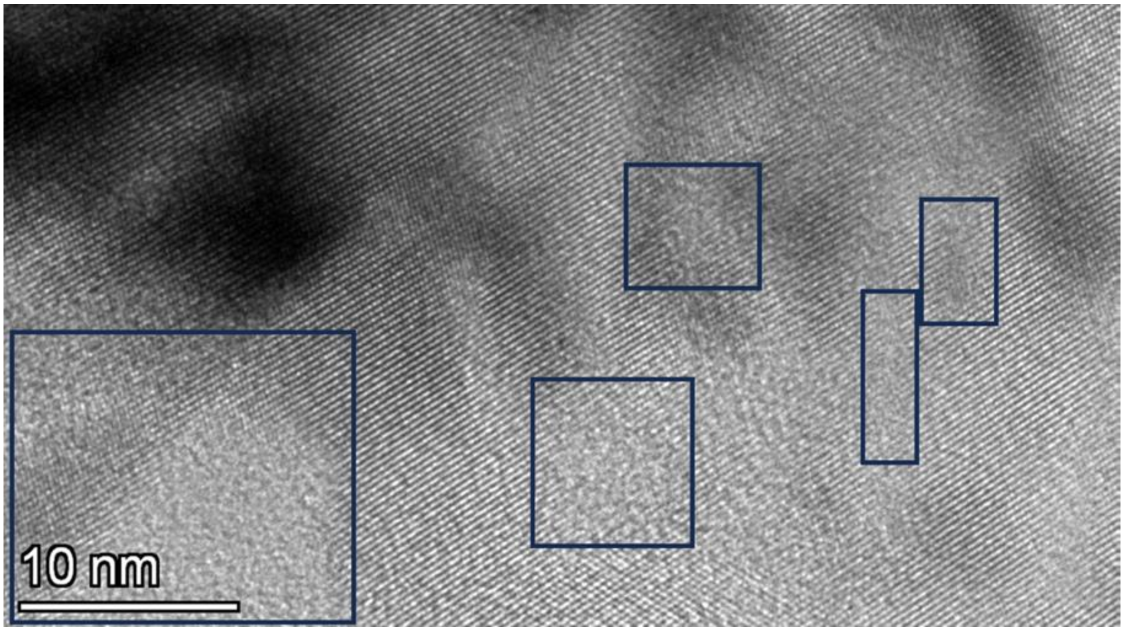


**Supporting Figure 7. HRTEM image showing amorphous regions (blue squared areas) within the orderly lattice fringes of *S. clausii* bacterial calcite.**

The Swiss cheese-like structures (Supporting Figure 8) has been thoroughly studied in other biomineral (21) and biomimetic (22) systems and have been shown multiple times to involve IOM. For example, a HAADF-STEM study of agarose-calcite composites showed dark low Z-contrast agarose-rich inclusions in calcite (22). Similar features have been observed in biominerals of higher level pluricellular organisms like nacre tablets of *Pinctada maxima* (23) or *Atrina rigida* (21). In the latter study, a density image analysis was used to infer that the pore-like structure in the shell was not due to voids but no further characterization was performed. Using HRTEM and STEM-EELS, de Frutos et al., (2) recently showed that organic fractions are present within these low calcium regions, usually enveloping small calcite lumps making up mollusc shells. Also, they showed that these areas do contain Ca, although in low quantities, and ACC was present. Our EDS line scans indicate that the Ca concentration does not drop to zero within the low Z-contrast pore-like areas, despite being zero outside the crystal edge, which coincides with observations of de Frutos et al. (2). A drop in the intensity for Ca in the EDS scans is expected because ACC being hydrated is less dense than anhydrous crystalline calcium carbonates, yet the presence of N along Ca, points to the joint presence of ACC and organics within these nanoregions.

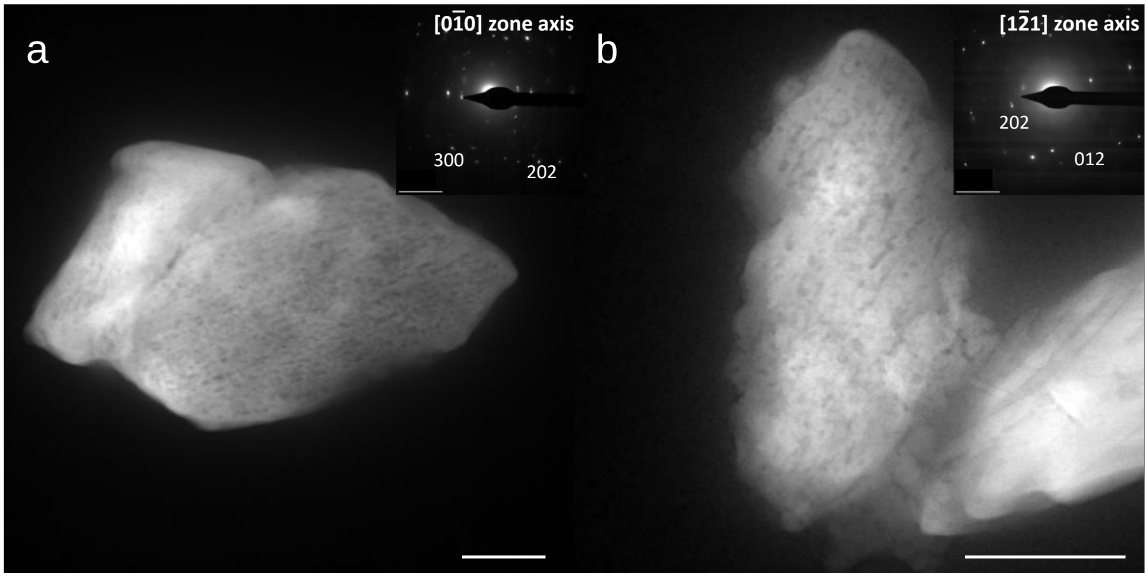


**Supporting Figure 8. TEM-HAADF images of bacterial calcites from *L. fusiformis* (a) and *S. clausii* (b) with predominantly parallel pore-like structures**. Inset: SAED patter of the crystals showing angular spreading. Scale bars 100 nm. Scale bars of SAED inset 5 $nm^{-1}$.

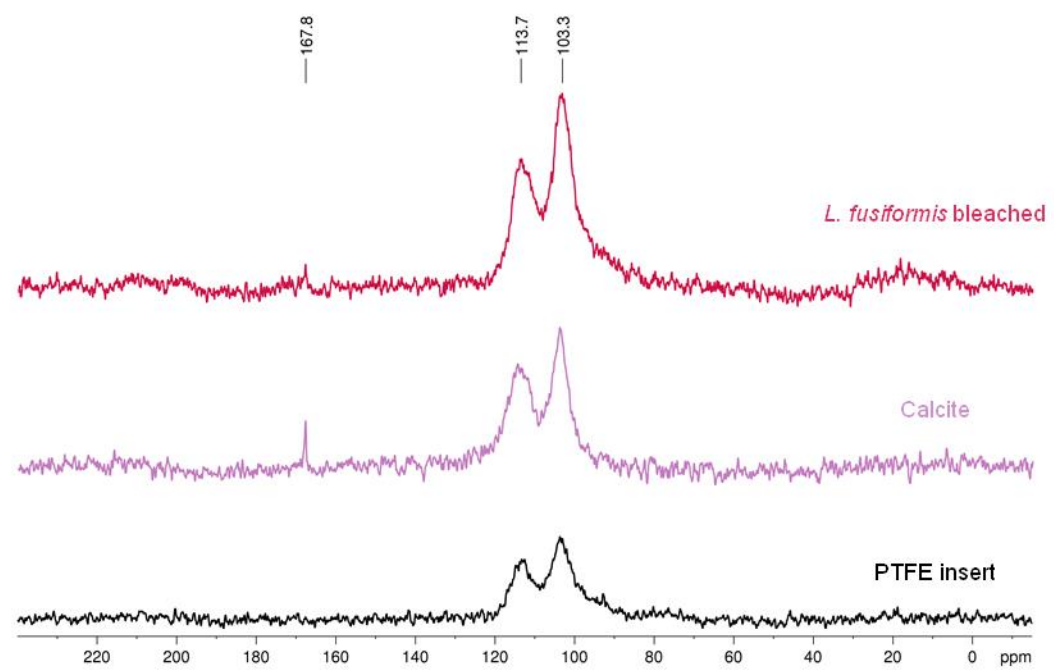


**Supporting Figure 9. MAS-NMR $^{13}C$ spectra of bleached *L. fusiformis* calcite (red), synthetic calcite (purple) and PTFE insert from the ssNMR setup (black).** MAS $^{13}C$ spectra of synthetic calcite shows a 167 ppm peak corresponding to the main carbonyl group and the peaks at 114 and 103 ppm from the PTFE insert. MAS-NMR $^{13}C$ *L. fusiformis* calcite shows a peak (of very low intensity) at 167.8 corresponding to calcite, and the same peaks at 114 and 103 ppm corresponding to the PTFE insert. All spectra are referenced to the signal of DSS ($\delta$ = 0 ppm).

**2D organic films**

Rodriguez-Navarro et al., (24) reported on how carbonic anhydrase produces β-sheet films that act as a blueprint on which ACC nucleates heterogeneously and then crystallizes as single crystal calcite. We presume that the bacterial organic 2D-crystalline film similarly acts as a template (

Supporting Figure 10 a, b) for the orderly formation of ACC previously stabilized with organic matter as mentioned above. These ACC particles act as small subunits which are finally crystallized into single crystal calcite with a shape not limited by (*hkl*) faces. The role of EPS in MICP is well described (25, 26) and observations have been done at the nanoscale of BCC on bacterial surfaces (27). Yet, *in vivo* reports of molecule-biomineralized calcite interactions during MICP are scarce. The closest report in literature found by the authors akin to the observed crystalline 2D structures were S-layers, although we cannot confirm those observed here are the same. S-layers are crystalline proteins, part of the cell envelope found in many types of bacteria, which reportedly can act as mineralization platforms (28–30). The S-layer shows protein units orderly placed at 3 to 5 nm (31) which match the *d*-spacings measured here. Furthermore, observation of fringes were also made in a similar fashion to those observed by us (29). Nevertheless, the current results preclude us from concluding that the structures observed here were S-layers, and other proteins are capable of self-assembling into 2D crystalline film structures (e.g. *79*). Nevertheless, this observation opens up the discussion of the diverse mechanisms of biomineralization in bacteria.

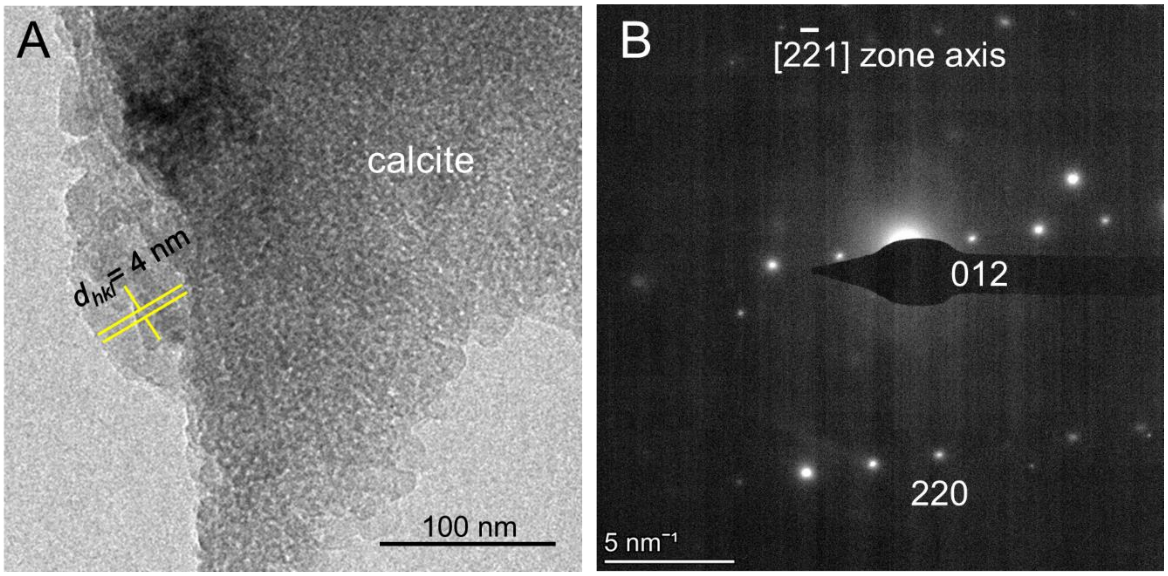


**Supporting Figure 10. (a) Crystalline 2D film showing lattice fringes (with d-spacing of 4 nm) with crystalline calcium carbonate formed on top from *L. fusiformis* precipitates. (b)** The corresponding SAED shows the single crystal features of calcite, which displays no crystalline faces and a nanogranular structure, consistent with its formation by aggregation of ACC nanoparticles followed by its template-directed crystallization on the 2D organic crystalline film. Note also that a very similar "frothy" structure was observed in calcite biominerals from bivalve shells, and according to Towe and Thompson (33) was due to intracrystalline occlusion of organics.
